# Novel cytometry-based characterization of lysosomal storage disease affected patient’s cells

**DOI:** 10.1101/2025.03.18.643908

**Authors:** Marine Laurent, Jérémie Cosette, Giulia Pavani, Sarah Bayol, Christine Jenny, Rim Harb, Julie Oustelandt, Anais Brassier, Daniel Stockholm, Mario Amendola

## Abstract

Wolman disease (WD) is a severe lysosomal storage disorder characterized by fatal lipid accumulation caused by the deficiency of a lipid metabolic enzyme, Lysosomal Acid Lipase (LAL), involved in the lysosomal hydrolysis of cholesterols and triglycerides. Due to the imbalance of lipids homeostasis, WD patients suffer from severe hepatosplenomegaly, hepatic failure and adrenal calcification resulting in a premature infant death within the first year of age. In this work, we explored multiple imaging analyses to fully characterize the phenotype of LAL deficient cells. In particular, we stained WD patients’ fibroblasts for intracellular lipid droplets (LD) and lysosomes and we analysed staining intensity and granularity as well as an increased number of LD and lysosomes using fluorescence wide field microscopy, confocal microscopy, conventional and image flow cytometry. Noteworthy, we showed that lipid homeostasis was restored upon delivery of a functional LAL transgene.

Finally, since fibroblasts cannot be used as routine clinical test as they are difficult to collect from WD patients, we confirmed our observations in LAL deficient human blood cell lines and in peripheral blood mononuclear cells (PBMC) from LAL deficient (LAL-D) mouse model, as a proxy for easily accessible WD PBMC.

Overall, we expect that this novel imaging analysis pipeline will help to diagnose WD, follow its progression and evaluate the success of enzyme replacement therapy or gene correction strategies for WD as well as other lysosomal storage disorders.

## Introduction

Lysosomal storage disorders (LSD) are inborn errors of metabolism caused by defective lysosome functioning and toxic metabolites accumulation in various organs and tissues. Lysosomal acid lipase deficiency (LAL-D) is an autosomal recessive LSD caused by mutations in the lipase A (*LIPA*) gene encoding for the LAL enzyme, the sole enzyme responsible for lysosomal hydrolysis of neutral lipids into cholesterol esters and triglycerides (Li and Zhang, 2019). The phenotypic spectrum of LAL-D depends on the residual enzymatic activity, and it ranges from the severe infantile-onset (Wolman disease, WD, OMIM 620151; prevalence ∼0.325–1.11 cases per million births), with <2% of remaining functional LAL, to milder childhood/adult-onset (cholesteryl ester storage disease, CESD, OMIM 278000; prevalence 3.13–4.86 cases per million births), with 2-12% of remaining functional LAL. WD patients suffer from hepatosplenomegaly and hepatic failure leading to premature infant death in the first year of life if left untreated (Balwani et al., 2023; Li and Zhang, 2019; Aguisanda et al., 2017a; Pericleous et al., 2017).

In physiological conditions, the excess of circulating neutral lipids is stored in cellular organelles called lipid droplets (LD), which are involved in diverse functions as cell survival, lipid cell homeostasis, and protein storage (Olzmann and Carvalho, 2019; Cohen, 2018; Fujimoto et al., 2008). In LAL-D patients, both lysosomes and LD pathologically accumulate in cells and affect whole organ function (Pericleous et al., 2017).

Enzyme replacement therapy (ERT), the only clinically available treatment for LAL-D, consists in the systemic injection of the recombinant human LAL enzyme, named Sebelipase alfa, to restore lipid homeostasis (Burton et al., 2022; Pastores and Hughes, 2020; Potter et al., 2021; Erwin, 2017; Burton et al., 2015). Early diagnosis is crucial for effective treatment and long-term survival. Liver biopsy is considered one of the most reliable methods for evaluating liver abnormalities; however, the risk of biopsy-related morbidity and mortality and the costs associated with this procedure limit its widespread application (Tommaso et al., 2018). Therefore, WD diagnosis commonly relies on the quantification of LAL enzymatic activity on dried blood spots (Lukacs et al., 2017; Hamilton et al., 2012) and the appearance of WD symptoms in the first week of life (Kohli et al., 2020; Reiner et al., 2014), eventually confirmed by radiologic examination (Foladi and Aien, 2021; Shenoy et al., 2014) and identification of the underlying *LIPA* mutations (Scott et al., 2013). However, simple biomarkers to follow disease progression and therapeutic treatment over time are still missing.

Here, we describe multiple imaging analyses to discriminate between WD patients and healthy donors (HD) cells and to evaluate the benefit of LAL gene replacement strategy into WD patients’ fibroblasts. We first investigated lipids and lysosomes accumulation in WD fibroblasts through fluorescence and confocal microscopy (Aguisanda et al., 2017b; Pham et al., 2005; Greenspan et al., 1985). Then, to increase the number of acquired events and the speed of the analysis, we analyzed difference in lipids storage and granularity between WD and HD fibroblasts using flow cytometry. In addition, to combine fluorescence microscopy with the statistical power of flow cytometry, WD and HD fibroblasts were analyzed with imaging flow cytometry (IFC), which record an image of each single cell and allows simultaneous quantification and counting of organelles, LD and lysosomes, at single cell level.

Finally, in view of a potential clinical application, we used our analysis workflow to investigate lipid deposits in engineered *LIPA* KO white blood cell lines, mimicking easily accessible peripheral blood mononuclear cells (PBMC), and in PBMCs from a LAL-D mouse model.

Overall, patients and healthy donor cells were effectively and reliably distinguished using lipid, lysosome, and granularity spot counting via IFC offering an appealing pipeline for patient’s diagnosis and treatment follow-up.

## Materials and methods

### Cell culture

Human primary fibroblasts were obtained from the NIGMS Human Genetic Cell Repository at the Coriell Institute for Medical Research: GM11851 and GM06124 (WD fibroblasts) and GM08333 and GM00041 (HD fibroblasts) (Table 1). Fibroblasts were maintained in Dulbecco’s Modified Eagle Medium (DMEM) medium supplemented with 2mM Glutamax (Gibco by Life Technology), 15% of FBS (FBS, BioWhittaker, Lonza) and 100U/ml of penicillin/Streptomycin (Gibco).

**Table 1:** Table with patient age and reference for Healthy (HD1 and HD2) and Wolman Disease (WD2 and WD3) fibroblasts.

| Sample | Reference at<br>Coriell Institut | Age of patient<br>(months) |
| --- | --- | --- |
| HD1 | GM00041 | 3 |
| HD2 | GM08333 | 5 |
| WD2 | GM06124 | 3 |
| WD3 | GM11851 | 4 |

K562 cells (ATCC® CCL-243, erythroleukemia cells) and Jurkat (ATCC® TIB-152, Lymphoblasts) were maintained in RPMI 1640 medium containing 2mM glutamine and supplemented with 10% fetal bovine serum (FBS, Lonza, Basel, Switzerland), 10mM HEPES, 1mM sodium pyruvate and penicillin and streptomycin (100U/ml each; Gibco, Waltham, MA, USA).

### LV vector cloning and production

The WT LIPA cDNA was cloned into a 3rd generation lentiviral vector (LV) backbone (Follenzi et al., 2000). LV were produced by transient transfection of 293T using third-generation packaging plasmid pMDLg/p.RRE, pK.REV, and pseudotyped with the vesicular stomatitis virus glycoprotein G (VSV-G) envelope (Amendola et al., 2005). LV were titrated in HCT116 cells and HIV-1 Gag p24 content was measured by ELISA (Perkin-Elmer) according to manufacturer’s instructions. The final product was formulated in sterile X-vivo 20 (Lonza, Basel, Switzerland) and stored at −80 °C.

### Fibroblasts transduction

Fibroblasts were seeded and cultured in a 6-well plate (Nunclon Delta Surface, Thermofisher). Cells were transduced, 1 day after seeding, at a MOI of 100 with a VSG-pseudotyped lentiviral vector encoding the human LIPA gene.

### LIPA KO cell lines

#### Editing

Three LIPA g(uide)RNAs (Gene Knockout kit V2, Synthego, CA, USA) were diluted following manufacturer’s instruction and ribonucleoprotein (RNP) complexes were formed with 30 pmol of SpCas9 (ratio1:2), kindly provided by Dr. J.P. Concordet.

Jurkat and K562 cells were electroporated with 3 µL RNP, 30 μM of Alt-R Cas9 Electroporation Enhancer (IDT, Coralville, IA, US) and 18 µL of Amaxa buffer solution: SE cell line 4D-Nucleofector kit (Electroporation program: FF-120) and SF cell line 4D-Nucleofector kit (Electroporation program: FF-120) respectively according to manufacturer’s instructions (Lonza, Basel, Switzerland) (Pavani et al., 2020). The day after electroporation cells were washed once in sterile PBS and resuspended in 500 µL of fresh medium.

#### Quantification of the KO

Genomic DNA was extracted with QuickExtract™ DNA Extraction Solution (Lucigen, Middelton, WI, USA). A PCR amplicon was designed to cover the three gRNA cutting sites (Forwad primer: TCACTGCTACCTTGCCAGTG; Reverse primer: CCTTACTGCTAGTCATTAAATTCCTAGC) using KAPA2G Fast ReadyMix (Kapa Biosystem, Wilmington, MA, USA). PCR products were loaded in a 1% agarose gel. Since simultaneous gRNA cutting can result in genomic deletion, we observed 3 different amplicon sizes: 571bp (no deletion:WT sequence or single gRNA editing, Band 1), 500 bp (deletion: between gRNA 1&2 or gRNA 2&3, Band 2) 432 bp (deletion: between gRNA 1&3, Band 3) (Fig 4.D).

Each band was quantified using ImageJ and the % was calculated with the following formula:

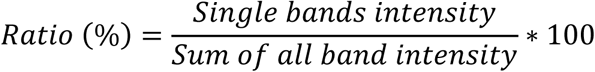

To calculate the % of indels in the undeleted band (571 bps), we gel purified the band, recovered the DNA, performed Sanger sequencing and TIDE analysis.

The LIPA KO score was calculated as follow:

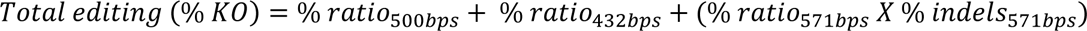

### PBMCs analysis on LAL-D mouse model

#### LIPA KO mouse model

The LIPA KO mouse model has a deletion of the exon 4 in the murine LIPA gene and was kindly provided by Pr. Kratky (Laurent et al., 2024; Pajed et al., 2019). Mice were maintained in a specific pathogen-free (SPF) environment with a regular light-dark (12 h/12 h) cycle and ad libitum access to food and water. This study was approved by ethical committee CEEA-51 and conducted according to French and European legislation on animal experimentation (APAFiS#24477-201912230940416 v1). 3-months LIPA KO and WT littermates were used for the blood processing.

#### Blood processing

Blood of LIPA KO and WT mice was withdrawn after isoflurane anaesthesia (IsoFlurin, Axience) from the retro-orbital veins with heparinized capillary and collected into 3,8% citrate Eppendorf tube. Then, the whole blood (around 100µL) was stained with Nile Red (NR) or LysoGreen (LG) (dilution 1:500, Abcam, Cambridge, UK) for 30 min at 37°C. After staining, blood was washed once with sterile PBS (Gibco by Life Technology) and centrifuged at 400 rcf for 2 min. The pellet was resuspended in red blood cells lysing and fixative solution (VersaLyse; Beckman Coulter, Pasadena, CA, USA) supplemented with 2.5% of IOTest 3 (Beckman Coulter, Pasadena, CA, USA) and incubated 10 min at room temperature. Finally, PBMCs were washed once in sterile PBS and resuspended in PBS solution prior flow cytometry and image flow cytometry analyses.

### Biochemical analyses

#### Western blot

To detect intracellular proteins, cells were lysed in RIPA buffer (Sigma-Aldrich, St.Louis, MI, USA) supplemented with protease inhibitor (Roche, Basel, Switzerland), freezed/thawed and centrifuged 10 minutes at 14,000 rcf at 4 °C. Total protein extract was quantified using BCA assay (Thermo-Fisher Scientific, Waltham, MA, US). 40 μg of protein were denatured at 90 °C for 10 minutes, run under reducing conditions on a 4–12% Bis-tris gel and transferred to a nitrocellulose membrane using iBlot2 system (Invitrogen, Waltham, MA, US). After transfer, membranes were blocked overnight at 4°C with Odyssey blocking buffer (Odyssey Blocking buffer (PBS), Li-Cor Biosciences, Lincoln, NE, USA) and incubated for 1 h with primary antibodies: mouse anti-hLAL (Thermofisher, 9G7F12, 7G6D7, dilution 1:2000) followed by specific secondary antibodies in PBS : Blocking buffer : goat anti-mouse IgG (Li-Cor Biosciences, IRDye® 800CW, dilution 1:15 000). α-actinin (sc-17829, mouse monoclonal, Santa Cruz Biotechnology, dilution 1:1000) was used as loading control. Blots were imaged at 169 μm with Odyssey imager and analysed with ImageStudio Lite software (Li-Cor Biosciences, Lincoln, NE, USA). After image background subtraction (average method, top/bottom), LAL band intensities were quantified and normalized with α-actinin signal.

#### LAL activity assay

Protein activity was detected in supernatants as previously described (Pavani et al., 2020; Aguisanda et al., 2017b; Hamilton et al., 2012) with minor modifications. Samples were incubated 10 minutes at 37 °C with 42 μM Lalistat-2 (Sigma-Aldrich, St. Louis, MI, USA), a specific competitive inhibitor of LAL, or with water. Samples were then transferred to a Optiplate 96 F plate (PerkinElmer), where fluorimetric reactions were initiated with 75 μl of substrate buffer (340 μM of 4-Methylumbelliferyl phosphate (4-MUP), 0.9% Triton X-100 and 220 μM cardiolipin in 135mM acetate buffer pH 4.0). After 10 minutes, fluorescence was recorded (35 cycles, 30 seconds intervals, 37 °C) using SPARK TECAN Reader (Tecan, Austria). Kinetic parameters (average rate, RFU/min) were calculated using Magellan Software and the relative LAL activity was calculated as follow:

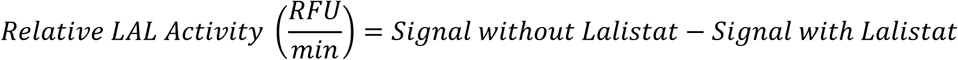

### Flow cytometry

Three hundred thousand Fibroblasts were detached with TrypLE (Gibco, Waltham, MA, USA) while 3 x 105 K562 and Jurkat cells were spun down (400 rcf 5 min) and stained with Nile Red (dilution 1:1000, Nile Red staining kit, Abcam, Cambridge, UK) or LysoGreen (dilution 1:1000, Lysosomal staining kit-Green Fluorescence, Abcam, Cambridge, UK) as per manufacturer’s instructions. Cells were analyzed using CytoFLEX S after laser calibration with Daily QC Fluorospheres (Beckman Coulter, Pasadena, CA, USA). Live cells were gated using forward and side scatter and ∼ 9 000 cells per sample were analysed. Unstained cells were used as negative controls. Data were analysed with Kaluza (Beckman Coulter, Pasadena, CA, USA).

### Image Flow Cytometry

Stained cells, as above, were acquired using an Image Stream X MKII (Amnis, Luminex, USA) with INSPIRE software (Amnis, Luminex, USA). Analyses were performed using IDEAS 6.1 software (Amnis, Luminex, USA). A first histogram of the gradient root mean square (RMS) of the bright field channel was plotted to exclude cells out of focus. A common Area vs Aspect Ratio dot plot allowed us to exclude cell doublets. Then, in the population of single cells, we analysed four parameters: the staining (NR or LG) fluorescence intensity and spot count per cell, the SSC (side scatter) autofluorescence intensity and spot count per cell. For SSC and NR, a spot is detected when its mean intensity reaches 20 times the average cell background ratio. For LG, a spot is detected when its mean intensity reaches 15 times the average cell background ratio. On average, 5 000 cells were analysed per experiment.

### Fluorescence Microscopy

Eighty thousand fibroblasts were seeded and cultured in a 6-well plate (Nunc, Thermofisher Scientific, Waltham, MA, US). After 5 days, cells were stained with Nile Red (Dilution 1:1000, Nile Red staining kit, Abcam) or with Lysogreen (Dilution 1:700, Lysosomal staining kit-Green Fluorescence, Abcam) as per manufacturer’s instructions. Eight fields for each condition were randomly acquired with an inverted fluorescence microscope (EVOS, Life Technologies, USA - 10X magnification) and average fluorescence intensity per cell was calculated with a custom-made ImageJ plugin. Images were first segmented using a manual threshold, which is chosen to optimize cell contour detection. Once thresholded, an area corresponding to the cell is automatically detected and the mean fluorescence intensity inside the cell is measured using the Analyze Particle feature.

### Confocal Microscopy

Thirty thousand fibroblasts were seeded and cultured in a MATEK (Ref: P35G-1.5-14-C, Mattek, Ashland, MA, USA). After 3 days, cells were first stained with Lysogreen (20 min at 37°C), washed with Dulbecco’s phosphate-buffered saline (DPBS) (Gibco, Thermofisher Scientific, Waltham, MA, US), stained for 15 min at 37°C with 5 µM Draq5 (ab108410, Abcam, Cambridge, UK), 10 µM Wheat Germ Agglutinin (WGA, W849, Thermofisher Scientific, Waltham, MA, US) and finally washed in Hanks’ Balanced Salt Solution (HBSS, Gibco). Plates were acquired with a LEICA TCS SP8 confocal microscope (LEICA, Germany) using a HC PL APO CS2 63X 1.4NA oil immersion objective. For each field of view, a stack of the whole cell layer was acquired (1 image every 0.4 µm). Several fields were acquired and quantified per condition. Three-Dimensional images of fibroblasts were reconstructed and quantified using Imaris Sofware (Oxford Instruments, UK). The total number of spots was measured using the spot feature of Imaris, with an estimated spot diameter set at 400 nm. The total number of spots was divided by the number of cells in the field of view, counted as the number of nuclei, to finally obtain the average number of spots per cell.

### Statistical analysis

Statistical analyses were performed using GraphPad Prism version 9.00 for Windows (GraphPad Software, La Jolla, CA, USA, “www.graphpad.com”). One-way analysis of variance (ANOVA) using Krustal-Wallis test were performed as indicated in the figures (alpha = 0.05, * p<0.032; ** p<0.0021; *** p<0.0002). Values are expressed as mean ± standard deviation (SD) as otherwise indicated. Only significant values are displayed in the figures.

## Results

We obtained and characterized primary fibroblasts from two healthy donors (HD) and two WD patients (Table 1). First, we confirmed that LAL protein and enzymatic activity were strongly reduced in WD compared to HD cells (Fig 1.A-B). Then, we analyzed the phenotype of fibroblasts by fluorescence microscopy. Cells were stained for neutral lipids or lysosomes, using Nile Red (NR) or LysoGreen (LG) dyes respectively. NR signal in WD fibroblasts was more intense than HD fibroblasts (∼1.4-fold; HD 5.7 ± 0.6 and WD 8.2 ± 1.4, mean ± SD), which confirms lipid accumulation in WD cells (Fig 1.C-D). A similar result was observed for LG (HD 7.9 ± 2.3 and WD 10.2 ± 2.1; mean ± SD) (Fig 1.C-E). In addition, using confocal microscopy and 3D images reconstruction, we could visualize (Fig 1.F) and count each lysosomal vesicle, and we observed that WD fibroblasts had ∼ 2.6 more lysosomes (Fig 1. G, HD 149.2 ± 27.3 and WD 391 ± 86.5; mean ± SD).

**Figure 1:**
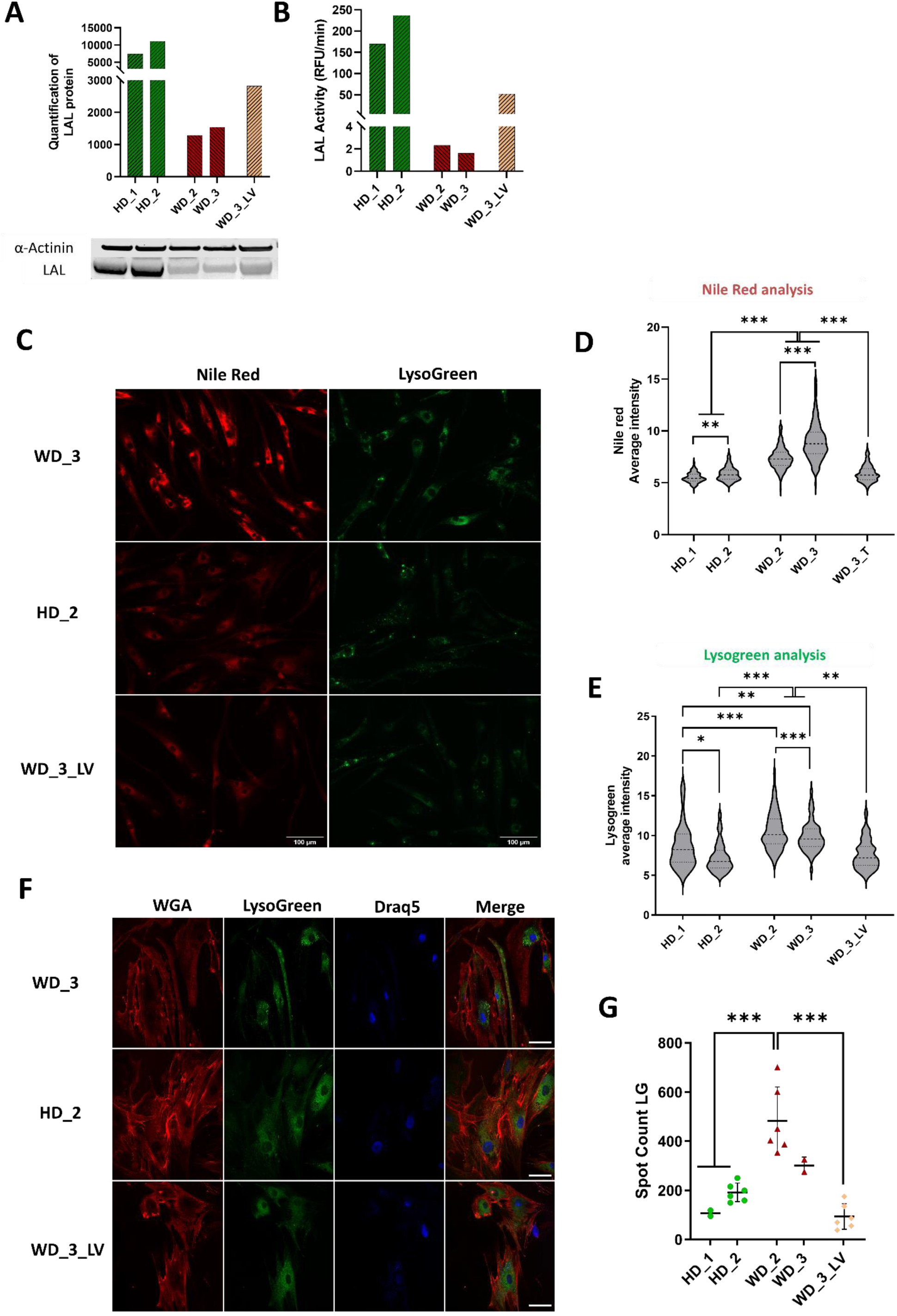
Biochemical and microscopic characterization of WD patients’ fibroblasts. **A**: Western blot of LAL and α-actinin proteins (lower panel) and quantification of LAL band normalized for α-actinin bands represented as fold over WD_3 (upper panel). **B**: LAL relative enzymatic activity represented as fold over WD_3. **C**: Fluorescence microscopy images of fibroblasts stained with Nile Red (left panel) or LysoGreen (right panel). Scale bar of 100 µm is in white. **D-E**: Quantification of mean fluorescence intensity per cell of Nile Red (NR; D) (HD_1 = 270 cells, HD_2= 258 cells, WD_2 = 438 cells, WD_3 =288 cells, WD3_LV = 253 cells) and LysoGreen (LG; **E**) (HD_1 = 42 cells, HD_2 = 105 cells, WD_2 = 134 cells, WD_3 =113 cells, WD3_LV= 134 cells). Each dot represents one cell. Bars represent mean ± SD. Kruskal-Wallis Test *p< 0,033 **p< 0,002, ***p<0.0002. **F**: Confocal microscopy images of fibroblasts stained with Wheat Germ Agglutinin (WGA) for cell membranes, LysoGreen for lysosome, Draq5 for the nucleus, and merge. Scale bar of 50 µm is in white. **G**: Number of LysoGreen spots per cell based on 3D confocal microscopy images. Each dot represents the average of spots per cells normalized per nucleus and analysed in a field of view (z-sack). Total analysed cells: HD_1 = 21 cells, HD_2=87 cells, WD_2 = 94 cells, WD_3 =69 cells, WD2_Treated = 62 cells, WD3_LV = 18 cells. Bars represent mean ± SD. Kruskal-Wallis Test ***p<0.0002. Statistical comparisons were made between each group, and only the differences that were statistically significant are displayed.

To confirm that the WD fibroblast phenotype is due to the absence of LAL and can be corrected with gene therapy, we introduced a functional hLAL transgene using a lentiviral vector (WD3_LV). After transduction, fibroblasts showed an increase of LAL expression and activity (Fig 1.A-B) resulting in a significant reduction of lipid (Fig 1.C-D, NR intensity: 5.9 ± 0.8; mean ± SD), lysosome intensity (Fig 1.C-E, LG intensity: 7.4 ± 1.6; mean ± SD) and lysosome number (Fig 1.F-G, WD_LV 93.4 ± 51.8; mean ± SD) compared to the untreated WD fibroblasts. Although these fluorescence microscopy analyses allow discriminating WD vs HD fibroblasts, they are time-consuming in slide preparation, acquisition and analysis of images, thus limiting the number of cells that can be analyzed.

Therefore, we decided to test flow cytometry, the gold standard method to measure cell fluorescence signal in a high throughput manner (Hasegawa et al., 2013). Confirming microscope analysis, we observed higher lipid accumulation (NR intensity) in WD fibroblasts compared to HD (∼ 2.1± 0.6, mean ± SD) (Fig 2.A). A similar trend was observed also for LG (Fig 2.B). Since accumulation of lipids and lysosomes change the granularity of the cells (Lee et al., 2004; Marina et al., 2012), we analyzed the side scatter signal (SSC) parameter, which requires no cellular staining, and demonstrated that it is an additional valuable measurement to discriminate WD and HD fibroblasts (Fig 2.C, ratio WD/HD: 2.5±0.7; mean ± SD). Finally, we demonstrated that the insertion of a functional copy of *LIPA* transgene fully restored lipid imbalance (Fig 2.A-C). Overall, flow cytometry represents an easy and rapid system to analyse WD fibroblasts and discriminate them from HD.

**Figure 2:**
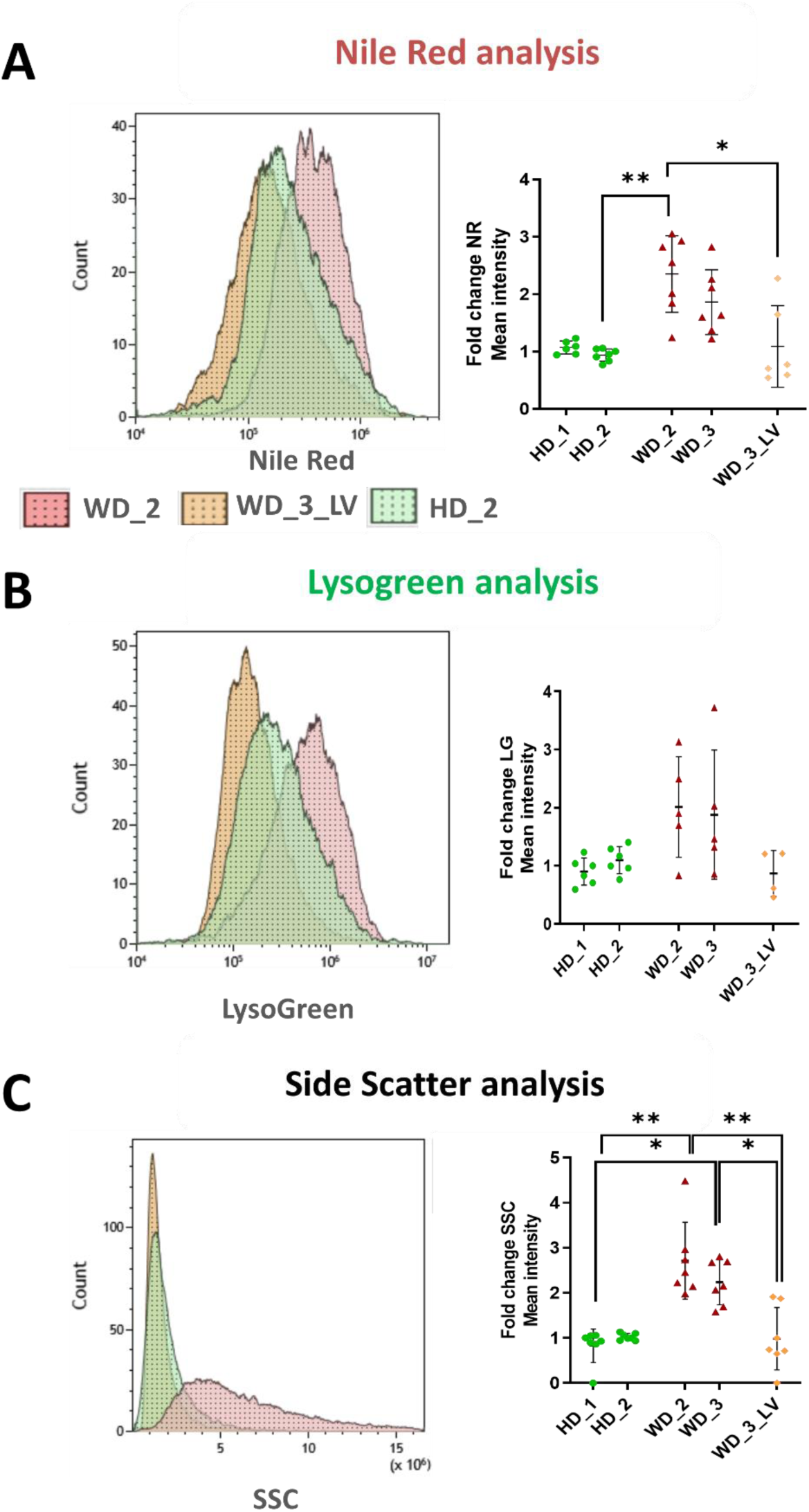
Conventional flow cytometry analysis of primary fibroblasts. **A**: Left panel: NR fluorescence intensity histogram; right panel: dot plot of NR intensity fold changes over HD_2. **B**: Left panel: LG fluorescence intensity histogram; right panel: dot plot of LG intensity fold changes over HD_2. **C**: Left panel: SSC intensity histogram; right panel: dot plot of SSC intensity fold changes over HD_2. Each dot represents an experimental replicate. Bars represent mean ± SD. Kruskal-Wallis Test; *p<0.033, **p<0.002. Statistical comparisons were made between each group, and only the differences that were statistically significant are displayed.

In order to combine the speed and statistic power of flow cytometry analysis with precise vesicle quantification offered by microscopy, we explored the use of IFC. Thank to single-cell imaging, IFC allows measuring fluorescence intensity as well as localizing and counting the fluorescent spots within each cell (Smirnov et al., 2017). In line with previous cytometry data, IFC showed a clear increase in NR signals in WD fibroblasts (Fig 3.A-B). Surprisingly, the counting of NR spots per cell allowed a better discrimination between HD and WD fibroblasts (∼400-fold of difference, Fig 3.C). Similar results were obtained when using SSC, where counting the spots of higher granularity was more efficient in discriminating HD and WD fibroblasts that measuring intensity (∼100-fold vs ∼ 4.5-fold; Fig 3D-E). For LG, instead, difference between samples was not clear either for intensity or for spot counting (Fig S1.A-C).

**Figure 3:**
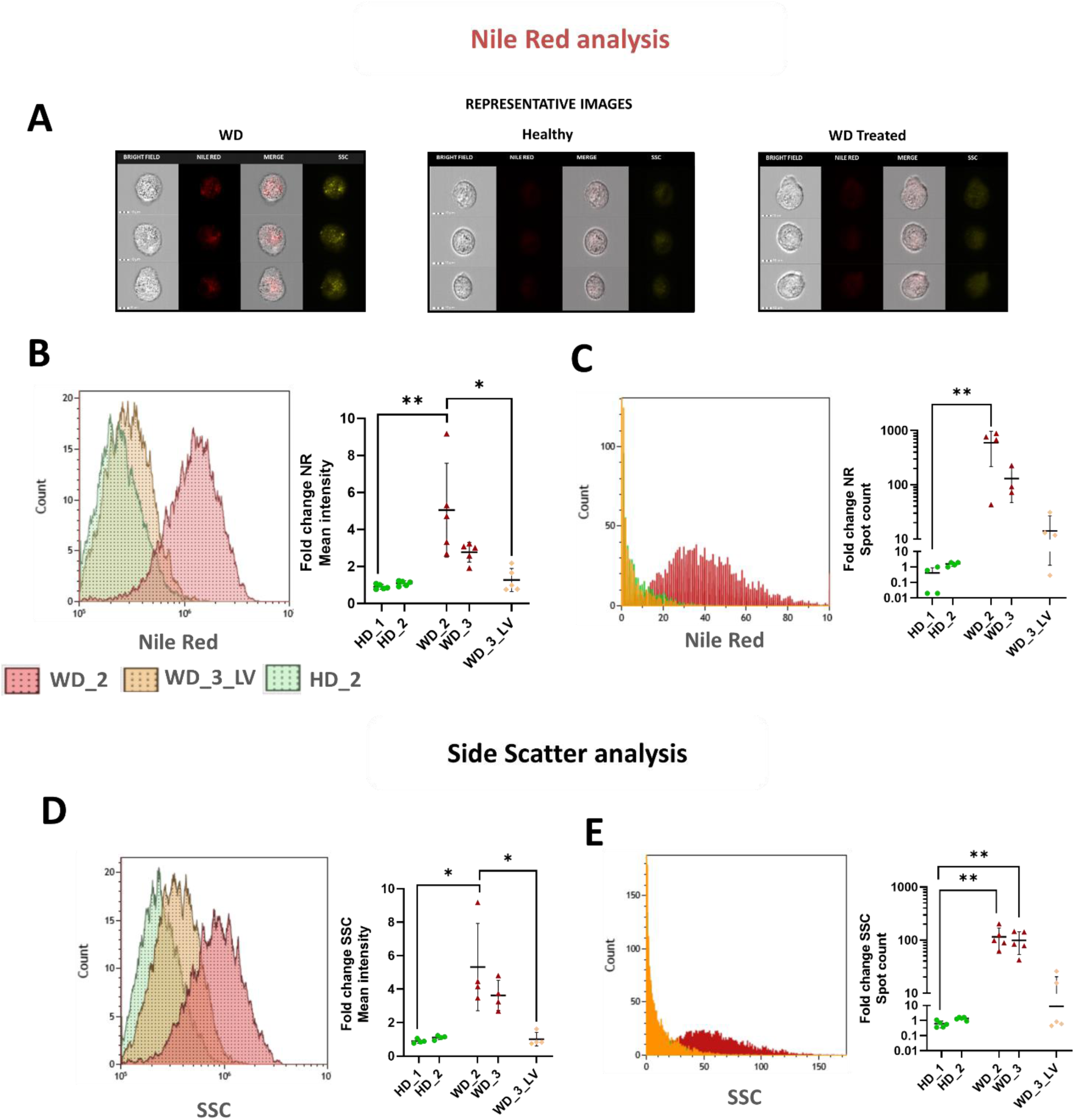
Imaging flow cytometry of primary fibroblasts. **A**: Representative image of NR-stained fibroblasts acquired by imaging flow cytometer. **B**: Left panel: NR fluorescence intensity histogram; right panel: dot plot of NR intensity fold change over HD_2. **C**: Left panel: histogram of NR spot count per cell; right panel: fold change over HD_2 of NR mean spot count per cell. **D**: Left panel: SSC signal histogram; right panel: dot plot of SSC signal fold changes over HD_2. **E**: Left panel: histogram of: SSC spot count per cell; right panel: fold change over HD_2 of SSC mean spot count per cell. Each dot represents an experimental replicate. Bars represent mean ± SD. Kruskal-Wallis Test *p<0.033, **p<0.002, ***p< 0.0002. Statistical comparisons were made between each group, and only the differences that were statistically significant are displayed.

Once more, we confirmed that the hLIPA cDNA encoding LV corrected cell phenotype and reduced the lipid accumulation (Fig 3.A-H).

Overall, spot counting via IFC seems to be a convenient and sensible method to evaluate lipid accumulation and discriminate WD vs HD fibroblasts, with SSC being a simple parameter for label free analysis.

Nevertheless, access to fibroblasts is difficult since it requires patient skin biopsies; therefore, we decided to test our analysis on blood cells. First, we validated our analysis pipeline using a LAL-D mouse model, with the deletion of exon 4 of the *LIPA* gene and lack of LAL activity (Fig 4.A), which mimics WD pathology (Laurent et al., 2024). PBMCs from WT and LAL-D mouse were analysed using NR, LG labelling or SSC signal with conventional flow cytometry or IFC. With conventional cytometry we observed a clear increase in lipid accumulation in LAL-D PBMCs in term of NR (ratio KO/WT: 3.7±0.6 fold, mean ± SD) and LG intensity (ratio KO/WT: 3.5±1.6, mean ± SD) and granularity (ratio KO/WT: 1.9±0.1, mean ± SD) (Fig 4.B). With IFC, we still observed a change in NR and LG intensity, but this difference was more pronounced for NR spot counting (ratio KO/WT: 6.8±1.1 fold, mean ± SD, Fig 4.C left panel) while no change was observed with LG spot count (Fig 4.C middle panel). No difference in granularity was observed between WT and LAL-D PBMC, in terms of both SSC intensity and spot count (Fig 4.C right panel).

**Figure 4:**
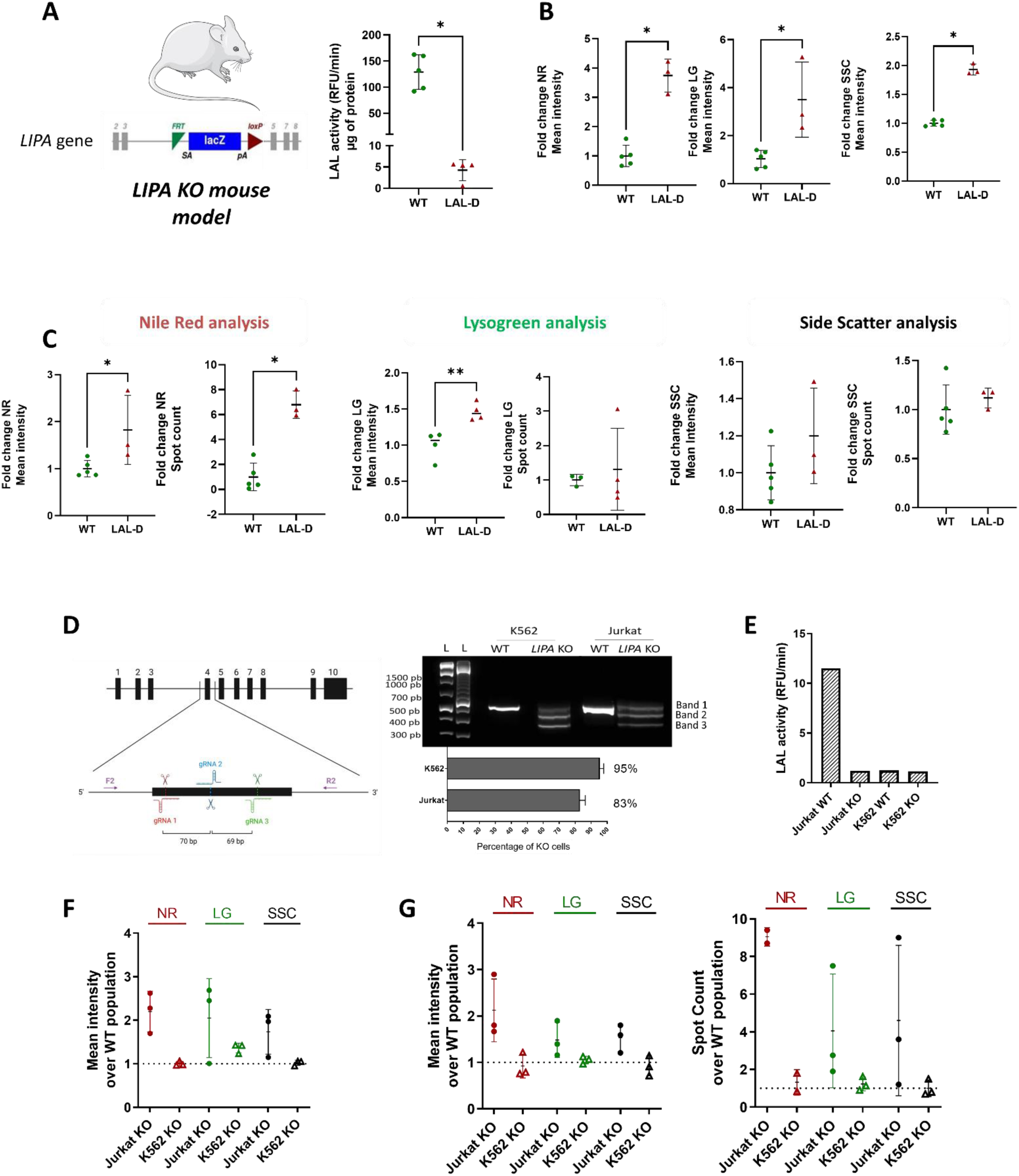
Characterization of LIPA KO mouse and human hematopoietic cells. **A:** Left panel: Representation of *LIPA* locus in LAL-D mouse with deletion of exon 4. Right panel: LAL activity in liver of 4 months-old LAL-D mouse. **B:** NR, LG and SSC MFI fold change over WT mouse acquired with flow cytometry. **C:** Left panel: NR mean intensity and spot count fold change over WT mouse acquired with imaging flow cytometer. Middle panel: LG mean intensity and spot count fold change over WT mouse acquired with imaging flow cytometer. Right panel: SSC mean intensity and spot count fold change over WT mouse acquired with imaging flow cytometer. **D:** Left panel: CRISPR/Cas9 editing strategy. *LIPA* gene structure is shown, with boxes indicating exons and lines indicating introns. Blow up is the exon 4 and surrounding genomic sequences. Arrows indicate PCR primers (F forward; R reverse); g(uide)RNA indicate the Cas9 cutting sites. The sizes of gRNA expected deletions (in nucleotides) are indicated underneath. Right panel: DNA agarose gel picture of PCR products of WT and edited cells. On the left, DNA size markers are shown. The histogram shows the percentage of *LIPA* KO. Bars represent mean ± SD. **E:** the LAL enzymatic activity in cell lines. **F:** NR, LG and SSC mean intensity fold change over WT cells acquired with conventional flow cytometer. **G:** NR, LG & SSC mean intensity and spot count fold change over WT cells acquired with imaging flow cytometer. Each dot represents one mouse or an experimental cell replicate. Bars represent mean ± SD. Kruskal-Wallis Test *p<0.033. Statistical comparisons were made between each group, and only the differences that were statistically significant are displayed.

Finally, we tested cytometric analyses on human blood cells. Since it is difficult to have access to peripheral blood mononuclear cells (PBMC) of patients, we generated *LIPA* KO blood cell lines using the CRISPR-Cas9 system. As for the mouse, we disrupted the exon 4 of the *LIPA* gene in erythroleukemia (K562) and T lymphocyte (Jurkat) cell lines by combining three different gRNAs (Fig 5.A) and we quantified the deletion efficiency via PCR (Fig 5.B) and the KO efficiency (Fig 5.C), by considering both the deletion and the indel efficiency. To confirm the *LIPA* KO phenotype, we measured LAL activity in cell pellets. While in Jurkat cells the editing completely abolished LAL enzymatic activity compared to untreated cells, in K562 cells no activity was detected in both WT and KO cells (Fig 5.D), suggesting that these cells do not express LAL and they were hence used as negative control. As described above, lipid and lysosomal vesicle accumulation was monitored using the NR and LG labelling or SSC signal with classical flow cytometry and IFC. Although all three parameters showed a similar increased signal in *LIPA* KO Jurkat cells with both cytometry methods (Fig 4.E-F), the spot counting with IFC provided the best discrimination between WT and KO Jurkat cells, with KO cells showing higher NR (∼ 9±0.5 fold; mean ± SD), LG and SSC (∼ 4 ± 3-fold; mean ± SD) (Fig 4.G). Finally, as expected due to the lack of LAL activity in WT K562, KO K562 cells did not exhibit any accumulation in lipid or lysosome vesicles (Fig 4.E-G), confirming the specificity of the imaging analysis.

In conclusion, IFC of easily accessible blood cells represents a fast and simple way to discriminate HD and WD phenotype quantifying NR or SSC intensity and spots number. We expect that this analysis will help in the diagnosis and clinical follow up of WD disease and its treatment, as well as of other inborn error of metabolism.

## Discussion

WD is a rare LSD leading to severe multi-organs accumulation of triglycerides and cholesteryl esters. In this study we explored the lipid deposits, lysosomal content, and granularity of WD patients’ fibroblasts with a combination of different imaging methods to improve diagnosis and clinical follow up of WD patients. With all imaging methods tested - fluorescence wide field microscopy, confocal microscopy, conventional flow cytometry and imaging flow cytometry - we confirmed accumulation of lipid vesicles in WD fibroblasts in comparison to healthy donor (Pham et al., 2005; Fukumoto and Fujimoto, 2002; Brown et al., 1988; Greenspan et al., 1985). In line with an increase of lipid content, we observed with both confocal and wide field microscopy a higher intensity of lysosome staining (LG) in WD cells (Aguisanda et al., 2017b), suggesting that lipid deposit increases the number or the size of lysosomes as a compensation mechanism for the loss of LAL function (Xu et al., 2012).

Although widely used to investigate the lipids content in WD, wide-field fluorescence microscopy (Aguisanda et al., 2017b; Pham et al., 2005; Brown et al., 1988) allows the processing and analysis of a limited number of cells and doesn’t allow a precise intracellular localization of the signal. In comparison, confocal microscopy provides 3-Dimensional reconstructed images of WD fibroblasts allowing to precisely count lysosomal vesicles and confirm their cytoplasmic localization (Fig 1.G-H). Nevertheless, confocal microscopy remains time-consuming and limited in terms of number of analyzed cells. To overcome these limitations, we analyzed WD fibroblasts via conventional flow cytometry and IFC (Fig 2-3). Both flow cytometry methods allow the analysis of a high number of cells in short time. Furthermore, together with fluorescence quantification, they allow measurement of the granularity change (SSC parameter) due to vesicle accumulation without any staining. With both methods we observed a clear increase in neutral lipids and granularity (SSC). IFC has the additional benefit of acquiring an image for each analyzed cell allowing localizing and quantifying the number of LD and lysosomal vesicles within the cell. This spot counting feature allows better discrimination of WD and healthy cells. In fact, we could observe a ∼100-fold increase of spots in WD fibroblasts based on NR staining or granularity (Fig 3.C-H). Since the number of LD increases more that their intensity, we can hypothesize that cells respond to the metabolic stress by generating more LD rather than enlarging them. To confirm that the phenotypic observation was due to impaired LAL function, we reintroduced in WD fibroblasts a functional copy of the *LIPA* gene. Corrected WD fibroblasts showed a significant reduction of lipid deposits and lysosomal vesicles confirming that LAL-D is the main cause of the phenotype, and that gene therapy is an option to restore lipids homeostasis (Fig 1-3).

Since access to WD patients’ material is limited by the rarity of the disease and to reduce interpatient variability, we investigated the lipid accumulation in murine PBMCs of LAL-D mouse model and in LIPA KO human hematopoietic cell lines. Interestingly, both the mouse and human cell models showed an accumulation of lipid vesicles and an increase of lipid content and granularity, confirming the feasibility of our proposed analysis pipeline on blood sample (Fig.4) for diagnosis and treatment follow-up.

This study has some limitations due to the dyes used for cellular staining. In fact, LDs are quite heterogeneous in nature and there is not a single dye that can identify all LDs. Therefore, a common strategy is to stain cells with fluorescent dyes with a strong affinity toward the neutral lipids of LDs, such as NR, to monitor and quantify LD in cells with the drawback of fluorescence background resulting from nonspecific binding to lipophilic sites other than LD (Smirnova et al., 2006). In addition, these dyes cannot provide information about the defective enzyme, resulting in the pathogenic condition. A similar issue affects LysoGreen, whose fluorescence intensity increases with pH acidification. Here, we have assumed that LysoGreen only stains lysosomes, however, while lysosomes are expected to exhibit the strongest fluorescence, other organelles, as Golgi apparatus, may contribute to the fluorescence signal (Demaurex, 2002; Marina et al., 2012). In addition, lysosomes are not only involved in lipid accumulation (Schultz et al., 2011), which could explain why LG signal is highly heterogeneous in cells (Fig 1.C-E) and does not always provide a clear distinction between WD and HD fibroblasts (Fig 3.E-F). Therefore, the combined use of NR, LG and SSC signals in term of intensity and spot counting should ameliorate the sensitivity and specificity of the analysis. Noteworthy, the specificity of the analysis is confirmed by the phenotypic amelioration observed with LAL gene replacement experiments.

We envision that the use of new more specific fluorescent dyes (Zhao et al., 2022) and antibody staining will improve cytometry analysis. Fluorescent dyes could also be combined with cell surface markers to which blood cell subpopulation is mostly affected by WD disease Finally, new IFC technologies (Schraivogel et al., 2022; Holzner et al., 2021) combined with artificial intelligence (Hirotsu et al., 2022; Otesteanu et al., 2021; Lippeveld et al., 2020) will allow faster image acquisition and more precise analysis as well as sorting and characterization of the cell of interest.

## Conclusion

In conclusion, we showed that lipid, lysosome, and granularity spot counting via IFC is a fast and sensitive method to discriminate between patient and healthy donor cells and to evaluate gene therapy treatment *in vitro*. Future tests on patients’ PBMC should elucidate the diagnostic and prognostic potential of this IFC analysis pipeline for WD as well as other lysosomal storage disorders.

## Acknowledgement

We thank Pr. Kratky Dagmar for providing the *Lipa^-/-^* mice (LAL-D) and Dr. Jean-Paul Concordet for providing Cas9 protein. We thank the whole Amendola’s laboratory for fruitful discussion. We also acknowledge Genethon “Vector Core Facility” for lentiviral vector productions and Genethon “Imaging and Cytometry Core Facility” for image analysis and Genethon “bioexperimentation team” for technical assistance on mice experimentation. We gratefully acknowledge the Conseil Général de l’Essonne (ASTRES) and Genopole Research in Evry for financial help for the purchase of equipment.

M.L. was partially supported by a PhD fellowship from the French Minister of Higher Education, Research and Innovation via University of Evry Val d’Essonne. M.A. acknowledges support by the Genethon, INSERM, the AFM-Telethon foundation (BE-DREP), the French National Research Agency (grants: NEEDED ANR -24-CE18-3712-01 ; PEMGeT ANR-22-CE17-0028-02; HemoLen ANR-20-CE17-0016-01; IRIS ANR-21-CE14-0063-03), the France Relance program, the DIM Thérapie Génique (SafeSCD), NextGenerationEU (PROGETTO CN00000041-SAFedit).

M.A. was also funded by the European Union (MAGIC grant 101080690, www.magic-horizon.eu; and EDISCD grant 101057659, https://editscd.eu/). Views and opinions expressed are however those of the author(s) only and do not necessarily reflect those of the European Union or HaDEA. Neither the European Union nor the granting authority can be held responsible for them.

