## Supplemental figure 1 for "Novel cytometry-based characterization of lysosomal storage disease affected patient’s cells"

**Table of supplemental figure:**

Supplemental figure 1: Imaging flow cytometry of primary fibroblasts. ..........................................1

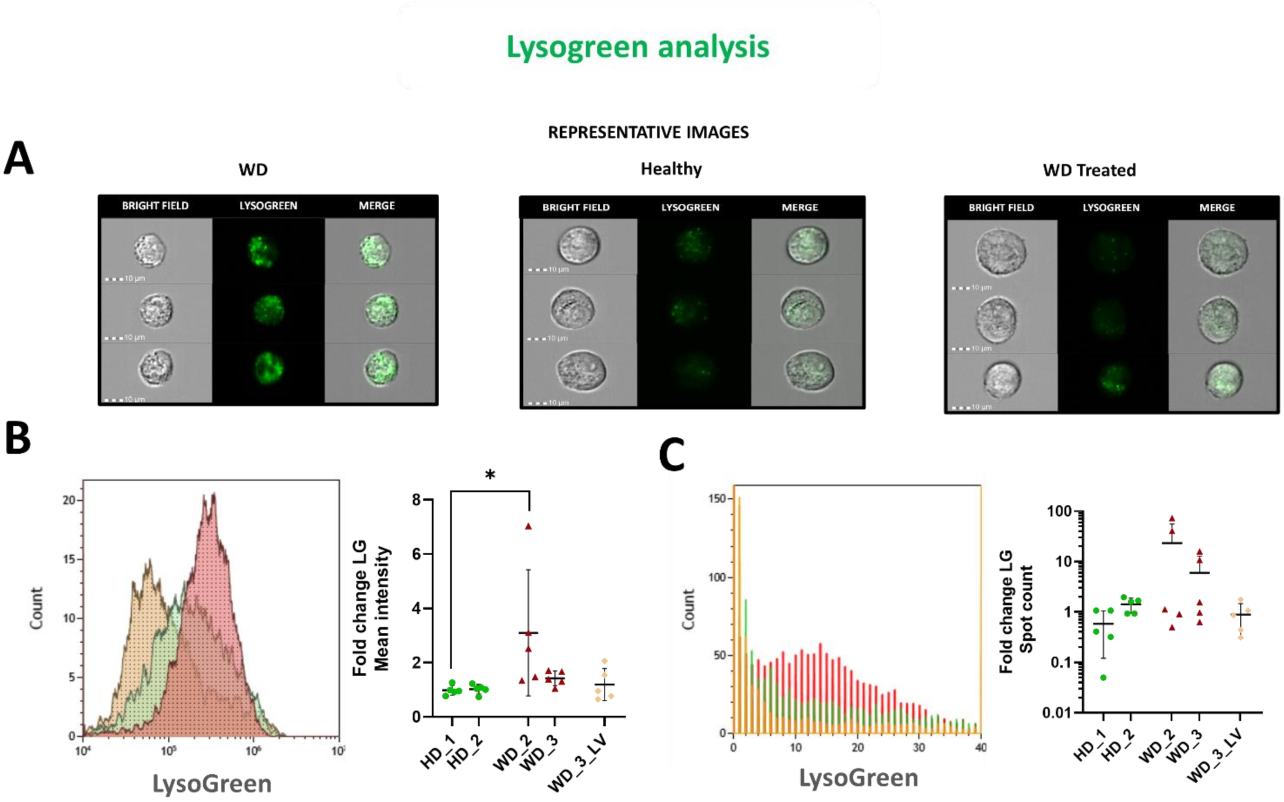


**Supplemental figure 1: Imaging flow cytometry of primary fibroblasts.**
**A**: Representative image of LG-stained fibroblasts acquired by imaging flow cytometer. **B**: Left panel: LG fluorescence intensity histogram; right panel: dot plot of LG intensity fold changes over HD_2. **C**: Left panel: histogram of LG spot count per cell; right panel: fold change over HD_2 of LG mean spot count per cell.
